# Assessing the impact of host density on vector abundance and transmission scaling of *Culicoides*-transmitted pathogens

**DOI:** 10.64898/2026.01.30.702916

**Authors:** Carly Barbera, Christie Mayo, Stacy Mowry, Jason R Rohr, T Alex Perkins

## Abstract

The spread of any pathogen depends on the dynamics of its hosts, with transmission rates typically assumed to either scale linearly with host population density (density-dependent transmission) or to be independent of it (frequency-dependent transmission). For vector-transmitted pathogens, a key determinant of transmission scaling is whether vector abundance is constant or sensitive to host abundance, with the latter being consistent with density-dependent transmission. Here, we assess whether *Culicoides* vector abundance increases in a manner that indicates density-dependent transmission of *Culicoides*-transmitted pathogens. To test this, we conducted trapping of *Culicoides* midges on eight livestock operations across a wide range of cattle abundances, placing traps at varying distances from the host aggregation. We used hierarchical Bayesian models to estimate the effect of host abundance on the vector-to-host ratio while accounting for differences in trapping efficacy between locations. Our results indicate a positive linear effect of host abundance on the ratio of vectors to hosts, with a posterior probability of 0.83. Median posterior values of the effect of host abundance on vector density predict a 2.3% increase (95% credible interval: −0.8, 23.2%) in the vector-to-host ratio with every 1,000 additional hosts. The weight of evidence from our study suggests that *Culicoides*-transmitted pathogens are likely subject to density-dependent transmission, and that transmission may be amplified in high-concentration livestock environments.

## Introduction

Pathogen spread is affected by numerous features of a pathogen’s environment, with the population ecology of the pathogen’s host being of primary importance. In particular, host density affects pathogen transmission through its influence on the rate of contact between susceptible and infectious hosts, which is a fundamental requirement for transmission to occur. Density-dependent transmission occurs in situations in which transmission is dependent on host population size (Keeling et al. 2008, McAllum et al. 2001, Lloyd-Smith et al. 2005). This is thought to be a plausible assumption for pathogens for which the transmission mechanism depends on the concurrent presence in a space by susceptible and infectious individuals, as is thought to be the case with measles, for example (Finkenstädt et al. 1998). In contrast, frequency-dependent transmission occurs in situations in which transmission is independent of host population size. This is classically thought to be a plausible assumption for vector-borne pathogens, which are transmitted by arthropods that take a limited number of blood-meals from hosts based on satiety, as is the case with malaria, for example (Anderson et al. 1991). In either case, the ecology of the pathogen and its host(s) shape the relationship between host density and the rate of transmission.

*Culicoides* biting midges are vectors for a number of arboviral diseases such as bluetongue, epizootic hemorrhagic disease, and Schmallenberg virus disease, that infect domestic and wild ruminant species, and for which the effect of host density on transmission is not clear. On the one hand, the viruses and their vectors circulate across landscapes with heterogeneous host population densities, potentially leading to increased contact rates in areas where there is a high degree of clustering of hosts (Keeling et al. 2008, McGregor et al. 2021, Charron et al. 2013). Livestock operations are highly variable in size, and are often adjacent to or overlapping with spaces occupied by wildlife species. This might lead to hotspots of transmission on large operations with high host densities, which would imply density-dependent transmission. However, this needs to be balanced with the fact that they are vector-borne pathogens. As a satiety-driven blood-feeding insect, the *Culicoides* midge vector may not produce any more contacts as hosts are added, which would imply frequency-dependent transmission. With the high degree of spatial heterogeneity in host populations, whether transmission is density- or frequency-dependent determines how well the transmission of a pathogen scales across a landscape of high-density commercial operations and smaller, more dispersed herds. This distinction can also have implications for control. If transmission is density-dependent, efforts to maintain host populations below a transmission threshold can be effective, but this does not apply if transmission is frequency-dependent (Keeling et al. 2008).

Currently, there is a lack of research that directly tests the effect of host abundance on vector abundance and, by extension, transmission of associated pathogens. There is previous work that suggests an effect, however. For example, Bakhshesh et al. (2020) found host density to be a significant risk factor for bluetongue virus (BTV) infection on the village level, and Calvete et al. (2009) found that cattle density and small ruminant farm density were positive predictors of BTV when modeling its distribution in Spain. Additionally, Jacquot et al. (2017) found livestock densities to play a key role in the spread of BTV across Europe, and Pioz et al. (2014) found a positive effect of beef cattle density on velocity of spread. However, other studies have found conflicting or negative results with regard to the effect of host density on infection prevalence. For example, both Chanda et al. (2019) and Nicolas et al. (2018) found varying effects of host density on BTV infection and velocity of spread, respectively, depending on the species of host examined. Pioz et al. (2012) found a negative effect of dairy cattle density on BTV spread, though they later found the opposite for beef cattle (Pioz et al. 2014). The lack of consensus points towards a need to explicitly test the effect of host abundance on vector abundance.

Theory predicts that the basic reproduction number, *R*_0_, is directly proportional to the number of vectors per host in a vector-borne pathogen system. *R*_0_ is defined as the number of infections generated by one infection in a fully susceptible population, which depends on the rate of contact between infectious and susceptible hosts (Dietz 1993, Hesterbeek et al. 2015). For vector-borne pathogens, contact rate is dependent on the ratio of vectors to hosts (Smith et al. 2004, Keeling et al. 2008), so assuming all other transmission parameters remain constant (biting rate, probability of transmission between vector and host during a bite, extrinsic incubation period, and host recovery rate), any increase in the ratio of vectors to hosts translates into a proportionate increase in *R*_0_. Thus, if this transmission system is density-dependent, we would expect to see an increase in the number of vectors per host as the number of hosts increases. In contrast, if the vector-to-host ratio does not increase with increased hosts, or reaches a threshold level at a very low number of hosts, this would indicate frequency-dependent transmission. Here, we expand on previous work by measuring changes in vector abundance at ecologically similar cattle sites spanning a wide range of host abundances. With this work, we can focus explicitly on the effect of varying host numbers on midge abundance.

## Methods

### Data Collection

We collected *Culicoides* midges on domestic cattle operations in Larimer and Weld Counties in Northern Colorado. Eight farms were selected, with cattle abundances ranging 200-69,000. Four were dairies and four were feedlots. To control for fine-scale habitat variation, four trap locations at each site were selected based on features common to all eight farms. At each farm, one trap was placed directly adjacent to a cattle pen, a crop field (corn at all sites), a road, and a water body or lagoon, for a total of four traps per farm. All sites contained a water body, referred to as a “lagoon,” though the size of the lagoons varied between sites. Figure 1A shows a schematic of a generic study site containing the habitat features at which traps were placed. While every “host trap” was the minimum distance away from the host aggregation, all other habitat types could vary in distance from the host aggregation based on the layout of that particular farm site. Varying the distance of the traps allowed us to control for any reduction in midge abundance when trapping farther away from hosts. The maximum distance from host pens was 400 m, based on the area of farms and accessibility of trapping locations. Traps were affixed to fence posts and were placed approximately 1.5 m above the ground.

**Figure 1.**
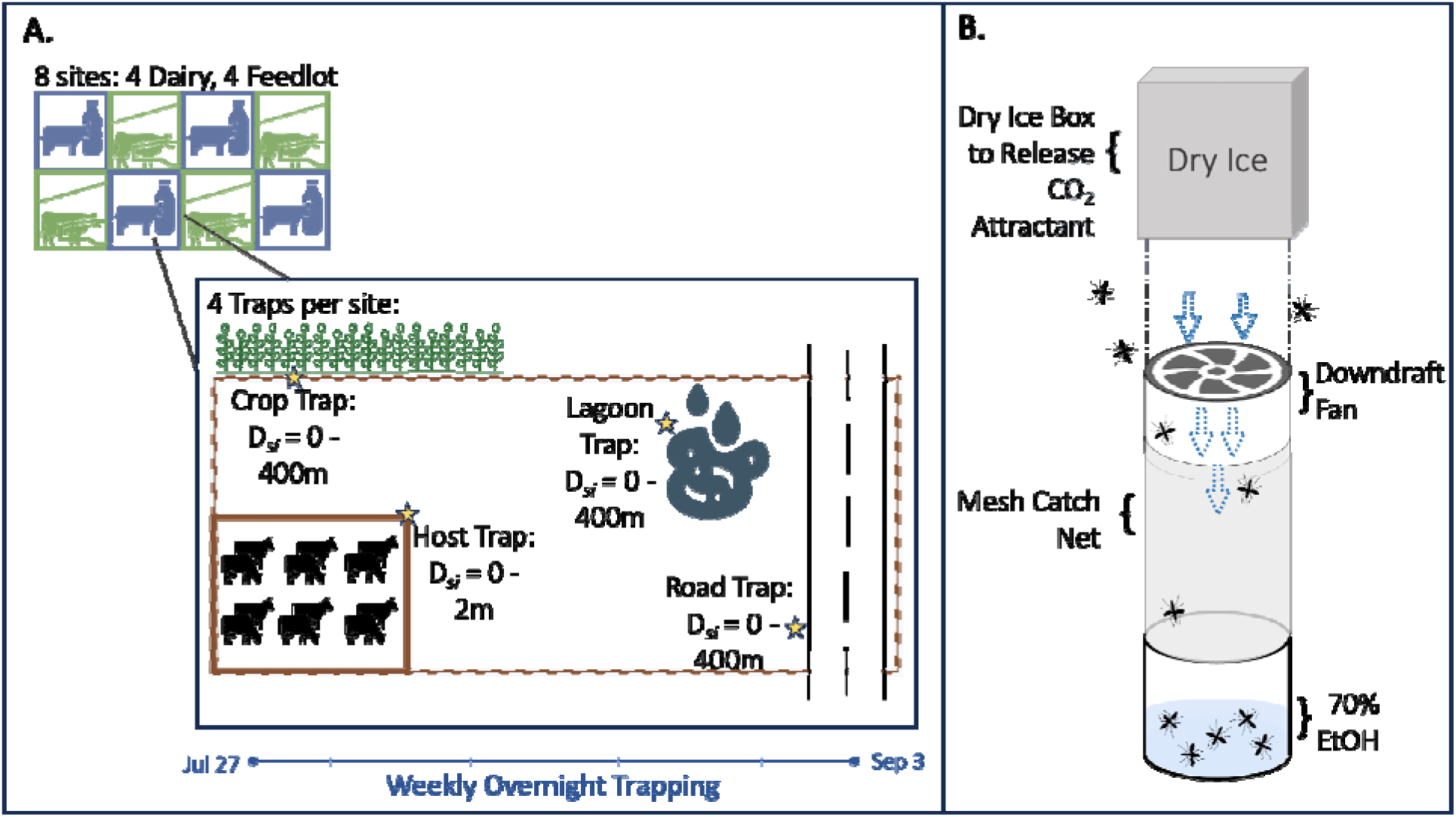
A) Schematic showing trapping set-up on a generic field site. Sites were either dairy operations or feedlots. Yellow stars indicate placement of a trap. Each site contained four traps, placed on one of each habitat type (host, crop, lagoon, and road). Host traps were, by definition, very near to cow pens. Other traps ranged in distance away from cow pens depending on where on that particular operation the specified habitat type was situated, with a maximum distance of 400 m. Trapping was conducted overnight once per week at each site. B) Trap design. Traps used dry ice, which releases CO_2_ upon sublimation, as bait to attract females looking for a bloodmeal, which can then be pulled into the trap by the downdraft created by the fan. Ethanol in the catch containers preserved captured specimens.

Midges were collected using CDC-style downdraft traps, baited with dry ice to release CO_2_ to mimic host attractant, with 70% EtOH to catch and preserve specimens (Figure 1B). Traps were not baited with light. Traps were assembled using vented foam boxes to hold and allow CO_2_ release from dry ice. Styrofoam boxes were connected with chains connected to a 4” diameter PVC tube with a 2.5” plastic propellor (Zoro brand Model G1630334) suspended inside. The propellor was secured to a small motor (Tasharina Corp. DC 6V 7000 RPM), which was connected to a 6V battery (Mighty Max 6V 4.5 AH rechargeable battery). A mesh sleeve was fitted on one end to the PVC downdraft piece and on the other to a plastic catch cup containing ∼200 mL of 70% EtOH. Trapping was conducted weekly at each site, with traps running overnight from approximately 7 pm to 8 am, to capture the timing of peak *Culicoides* activity at dawn and dusk. Specimens from each trap were stored in 70% EtOH at −20 °C until counting of midges from each sample. We obtained information on approximate numbers of cattle at each site from farm managers.

### Model formulation

To evaluate the effect of host density on per-host midge vector abundance, we used a model fitted to the midge abundance data with Bayesian inference. The model is visually represented in Figure 2, and a full list of variables and parameters is provided in Table S1.

**Figure 2.**
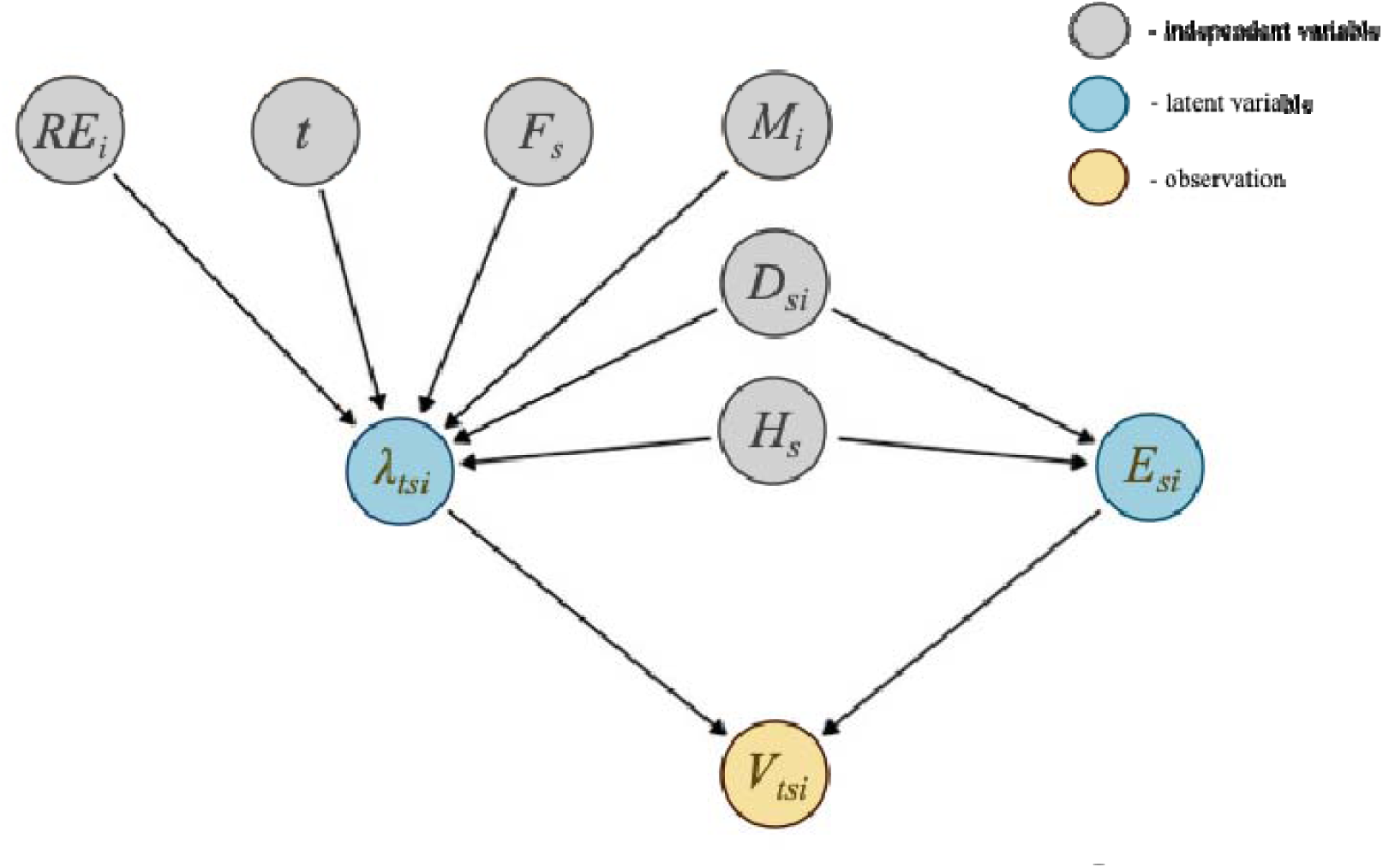
Model schematic for the most complex version of the model. The yellow circle contains *V_t,s,i_*, the observed captured midges at time *t*, site *s*, and trap location *i*. Blue circles contain the unobserved latent variables (λ*_t,s,i_* = true midge abundance, *E_s,i_* = trap efficiency, respectively). Grey circles contain independent variables (*RE_s,i_* = random effect of trap location, *t* = time point, *F_s_* = site type, *M_i_* = microhabitat type, *D_s,i_* = distance from hosts, and *H_s_* = host abundance). Arrows denote the directionality of effects of one variable on another.

We modeled observed midge vector abundance on day *t* at site *s* and trap location *i* as a negative binomial random variable with mean *V_t,s,i_*, such that

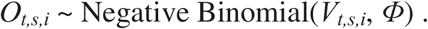

A negative binomial distribution is a common choice for modeling count data, especially when variability in counts is potentially high (Lindén et al. 2011). The variable *V_t,s,i_* is jointly dependent on two latent variables: trap efficiency, *E_s,i_*, and “true” midge abundance, λ*_t,s,i_*, such that

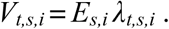

Estimating *E,_s,i_* allows us to account for reduced effectiveness of the CO_2_-baited traps when placed near large numbers of actual hosts, which may divert midges away from the traps with their own CO_2_. It is common to place traps a set distance apart from each other to reduce interference of one trap’s bait with another’s (Kirkeby et al. 2013, McDermott et al. 2016), so it follows that host cues could have a similar interfering effect. Indeed, other studies have noted that host animals themselves can be greater attractors of midges than baited traps are (Elbers et al. 2023, McGregor et al. 2019). Thus, by accounting for reduced trap efficiency due to proximity of hosts, we can separate the more ecologically relevant “true” midge abundance from captured midge abundance *V_t,s,i_*.

Trap efficiency was modeled as a function of host headcount *H_s_*, distance of trap from the host aggregation *D_s,i_*, and the interaction of the two (*H_s_D_s,i_*), as the negative effect of a high head count on trap efficiency was expected to be moderated when moving farther away from the host aggregation. Trap efficiency *E_s_*_,i_ is a probability and must be between 0 and 1, resulting in

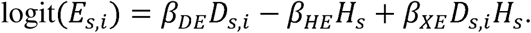

The variable, λ*_t,s,i_*, or ‘true’ midge abundance, was modeled as an effect of site type *F_s_*(dairy vs feedlot), distance *D_s,i_*, scaled head count *H_s_*, and time *t* in the trapping season. Site type was assumed to have an impact on midge abundance due to differences in land management on dairies compared to feedlots. For example, the use of agricultural wastewater lagoons to store cow manure on dairies potentially leads to differential availability of larval habitat. Distance from host aggregation *D_s,i_* was included to account for changes in midge abundance farther from hosts. Trap microhabitat type *M_i_* was included to account for any impact of within-site land feature differences on midge abundance at a given trap. We also included a random effect of site, *RE_s_*, to account for random differences in midge abundance between field sites not explained by covariates in our model. The equation is multiplied by HRaw_s_, or the unscaled head count, to model the midge abundances on an appropriate scale. To ensure midge abundance was strictly positive, we used a log link for λ*_t,s,i_*, resulting in:

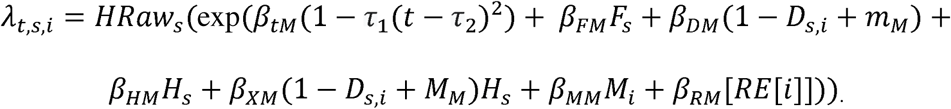

The expression 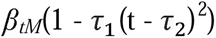 describes the seasonal curve of midge abundance, with parameter *β_tM_* describing the height of the curve, τ_1_ describing the width of the curve, or the difference in magnitude between peak abundance and minimum abundance, and τ_2_ describing the timing of the peak. Day in the season was scaled between 0 and 1, with the earliest day (July 27) as 0 and the latest day (September 3) at 1. The effect of trap distance *D_s,i_*on midge abundance was modeled as *β_DM_*(1 - *D_s,i_*), because we *a priori* expected midge abundance to decrease farther from hosts (Rigot et al. 2012), with parameter *M_M_* included to represent the proportion of midges at the furthest distance compared to the closest, and to prevent there from being zero midges at the maximum distance *D_s,i_* = 1. An interaction between *D_s,i_* and *H_s_* was included to capture any moderating effect of distance on the effect of head count on λ*_t,s,i_*.

### Model Fitting and Selection

The model presented above is the full version of the model that includes all variables of biological relevance for this analysis. We also considered alternative versions of the model that excluded covariates and interacting terms for which there is a possible, but unclear, biological basis. Specifically, we ran models that excluded the interaction between *D_s,i_* and *H_s_* as a covariate for the latent variable *E_s,i_*, for the latent variable λ*_t,s,i_*, or for both. Additionally, for each of these alternative models, we tested different ways of including covariates for the microhabitat type surrounding each trap. As described above, each site contained four microhabitat types, with the host trap type being, by definition, the minimum possible distance away from the host aggregation. Because of this, there is collinearity between microhabitat type and distance that could make it better to include only one of these variables. Thus, we ran models either excluding microhabitat entirely, including microhabitat as a binary variable (host trap type vs. non-host trap type), or including all covariates for four microhabitat types (i.e., host, crop, road, and lagoon), for a total of 12 models (Table 1). All models were also run without the trap-level random effect *RE_s,i_* to quantify how much of the variance explained by each model was driven by fixed versus random effects, and to be able to compare model performance among models with and without random effects, given that random effects can dramatically change model performance.

**Table 1.** Models organized by their inclusion of various interactions (rows) and inclusion of microhabitat variables (columns).

|  | **No Microhabitat Variables** | **Binary Microhabitat** | **All Microhabitat Variables** |
| --- | --- | --- | --- |
| **No Interactions** | Model 1 | Model 2 | Model 3 |
| **Interaction in E** | Model 4 | Model 5 | Model 6 |
| **Interaction in $\lambda$** | Model 7 | Model 8 | Model 9 |
| **Interaction in Both** | Model 10 | Model 11 | Model 12 |

To evaluate whether increased host abundance leads to an increased number of vectors per host, we focused on values of the *β_HM_* parameter, which represents the effect of head count on λ*_tsi_.* Additionally, we used the estimates of *β_HM_*for each model to calculate the predicted difference in *R*_0_ between the site with the highest head count and the site with the lowest head count. Because *R*_0_ is proportional to the ratio of vectors to hosts and *β_HM_* describes the impact of host abundance on vector abundance, it follows that

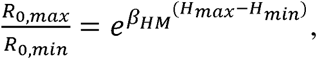

where *R*_0,max_ is the value of *R*_0_ at the site with the highest host head count *H*_max_ and *R*_0,min_ is the value of *R*_0_ at the site with the lowest host head count *H*_min_. Because *H_S_* is scaled between 0 and 1, and *H_max_* =1 and *H_min_* = 0, this simplifies to

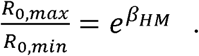

After the initial model comparison, which focused on models assuming a linear effect of host abundance on the vector-to-host ratio, we then compared the selected model to ones assuming a threshold for the effect of host abundance. Specifically, we ran two models assuming a type-II functional response (saturating function) and a type-III functional response (sigmoidal function), respectively (Holling 1959, Dawes et al. 2013). We then compared those models to the model with a linear function. If models assuming that host abundance only increases the vector-to-host ratio up to a point fit the data better than models assuming a linear increase, we could conclude that transmission is frequency-dependent beyond that point and the type-II and type-III models would replace the linear function of host abundance with 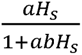 and 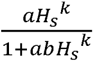, respectively.

We fitted models using Hamiltonian Markov chain Monte Carlo in ‘rstan’ version 2.21.8 in Rstudio (Stan Development Team 2025, Posit Team 2025). Chains were run for 10,000 iterations with a burn-in period of 5,000. Convergence of MCMC chains was assessed visually using the ‘bayesplot’ package (Gabry et al. 2019) and the Gelman-Rubin statistic. Model performance was evaluated using LOOIC estimates from the ‘loo’ package version 2.6.0 (Vehtari et al. 2020), and by calculating R^2^ from posterior predictions from each model. We calculated the R^2^ using the method proposed in Gelman et al. (2018), 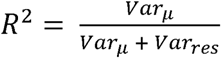, where *Var*μ is the variance of the modeled predictive means and *Var_res_* is the residual variance.

### Priors

Priors were chosen to be relatively uninformative, as many of our model’s parameters have not been well-quantified in the literature. Most parameters were assumed to be normally distributed with a mean of zero, consistent with a neutral prior understanding about whether effects of scaled predictor variables are positive or negative. Parameters τ_1_, Φ were constrained to be positive (gamma distribution with shape = 3 and inverse scale = 1), given that τ*_1_*describes the magnitude of the seasonal peak compared to the minimum and Φ is a variance. *β*_DE_ and *β*_HE_ also had prior gamma(3,1), to reflect the assumption that trap efficiency increased with increasing distance from hosts, and decreased with increasing host abundance. Parameters *M_M_*and τ*_2_* were constrained between 0 and 1 with a uniform prior. Full details of prior assumptions are summarized in Table S1.

## Results

### Data Collection

A total of 156 trap-nights were collected from the eight sites between July 27 and September 3, 2020. Each of the 32 unique trap locations was visited an average of 5.4 times (range: 3-7, as some samples were lost due to trap malfunction or other circumstances). In total, 2,241 midges were trapped, with abundance per trap-night ranging from 0 to 130 midges. However, average numbers of midges per trap-night was much lower (14.4), and many trap-nights (44) contained no midges. The timing of peak midge abundance within the season was estimated to be August 21-22 (τ_2_ Median: 0.65; 95% CrI: 0.58, 0.82) (Figure S1). The lowest midge abundance was estimated to occur at the earliest collection date. All models estimated similar seasonal curves (Table S2 M,N,O).

### Model Fit and Comparison

Hamiltonian Monte Carlo chains converged for all 12 models based on visual inspection of trace plots and R̂ < 1.005. Based on WAIC and R^2^ values, all models performed comparably well, with ΔLOOIC between models not exceeding 1.7 and R^2^ values ranging 0.747 - 0.754. Standard errors of the LOOIC values for all 12 models overlapped, suggesting that no model fit the data distinguishably better than any other model on the basis of LOOIC.

All of the 12 aforementioned models included trap-level random effects. To understand the extent to which model performance was driven by those random effects, we fitted 12 otherwise identical models without random effects. This showed that between 68% (Model 3, Model 6, Model 9, and Model 12) and 83% (Model 1 and Model 4) of variance in the data explained by each model overall was accounted for by the random effects. This means that between 17% (Model 1 and Model 4) and 32% (Models 3, 6, 9, and 12) was explained by the fixed effects included in the model, rather than unexplained differences among traps. Models including all four microhabitat types showed the highest amount of variance accounted for by fixed effects.

### Parameter Estimates

Examination of parameter estimates across models revealed some differences resulting from exclusion of microhabitat variables, as well as from inclusion of interaction terms. In summary, models excluding microhabitat estimated that increasing distance from host aggregation increases midge abundance. Given that this effect was seen only in the models that excluded microhabitat, we concluded that these models are unable to account for an effect which then is misattributed to distance effects. Hence, the positive effect of distance on midge abundance in these models is not justified. Once microhabitat variables were included, there was very little difference between outputs of the models with two microhabitats and the models with all four microhabitat types. Inclusion of an interaction between *D_s,i_* and *H_s_* in *E_s,i_* did change the effect of head count on trap efficiency at different distances, but inclusion of this interaction for λ*_t,s,i_* did not meaningfully change the relationship. Thus, in the absence of differences in WAIC or R^2^ to choose a “best” model, we chose to focus on Model 5, which includes two microhabitat types and an interaction in the *E_s,i_* term. Results for this model are presented in figures in the main text, and any differences in effects between this model and others is noted in the text. This model had R^2^ = 0.748, 21.1% of its variance was explained by fixed effects, and its predictions generally matched the data (Figure 3).

**Figure 3.**
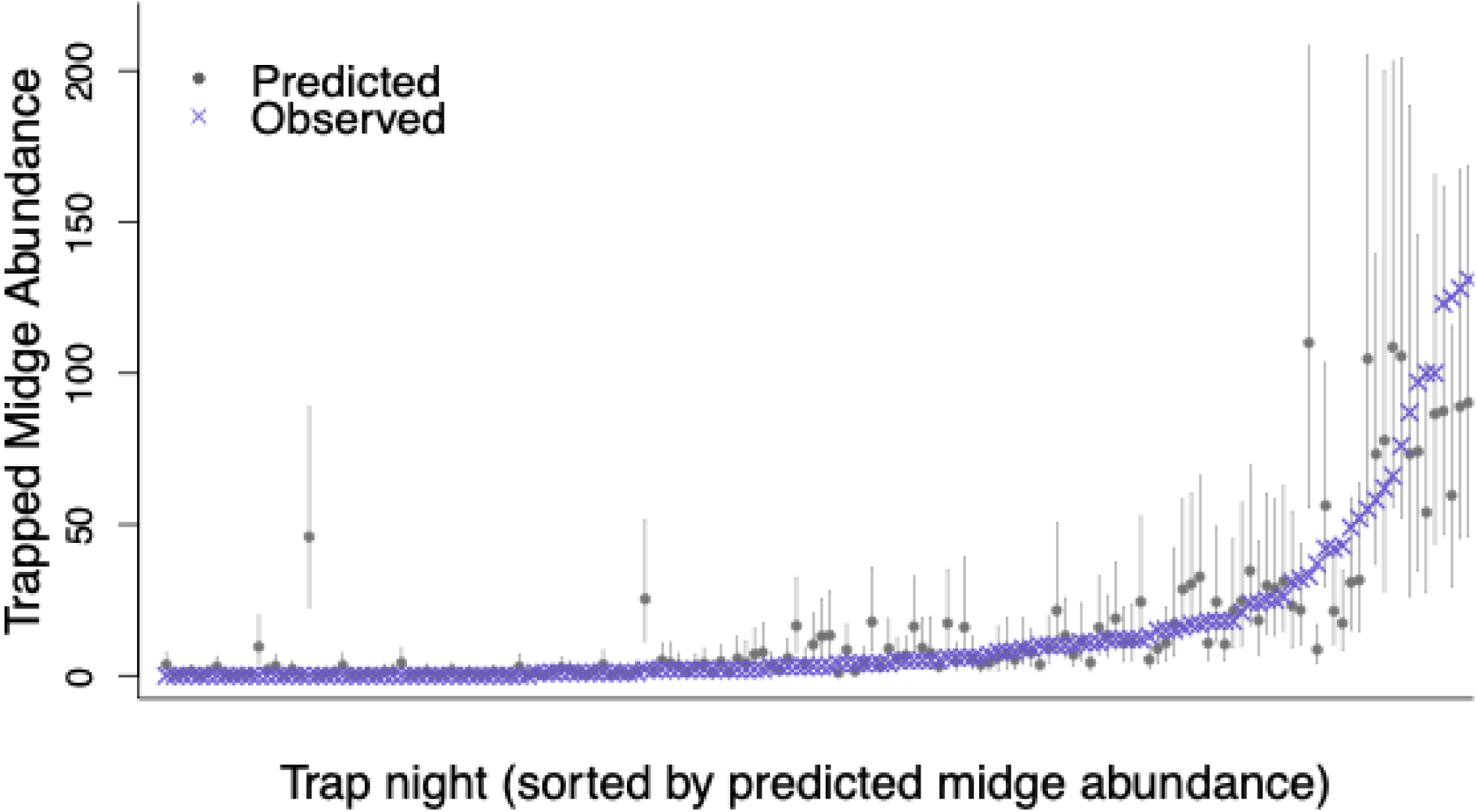
Posterior distributions of *V_t,s,i_,* trapped midge abundance, for Model 5. Gray circles and whiskers are the median and 95% posterior predictive intervals for each data point. Blue x marks indicate the observed values.

#### Trap Efficiency *E_si_*

In Model 5, trap efficiency was estimated to decrease with increasing host abundance (*β_HE_*median 2.14; 95% CrI: 0.56, 4.78) and increase with greater distance from host aggregation (*β_DE_* median 3.24; 95% CrI: 0.83, 8.26). Additionally, we estimated a positive interaction between distance and head count (*β_XE_* median: 0.74; 95% CrI: −2.87, 5.32), meaning that the effect of host abundance was weaker at further distances (Figure 4A, B) and, likewise, that the effect of distance was stronger at higher host abundances (Figure 4C, D). The direction of all of these effects are consistent with the hypothesis that hosts compete with traps for nearby midges due to the fact that midges are attracted to both based on their emission of CO_2_.

**Figure 4.**
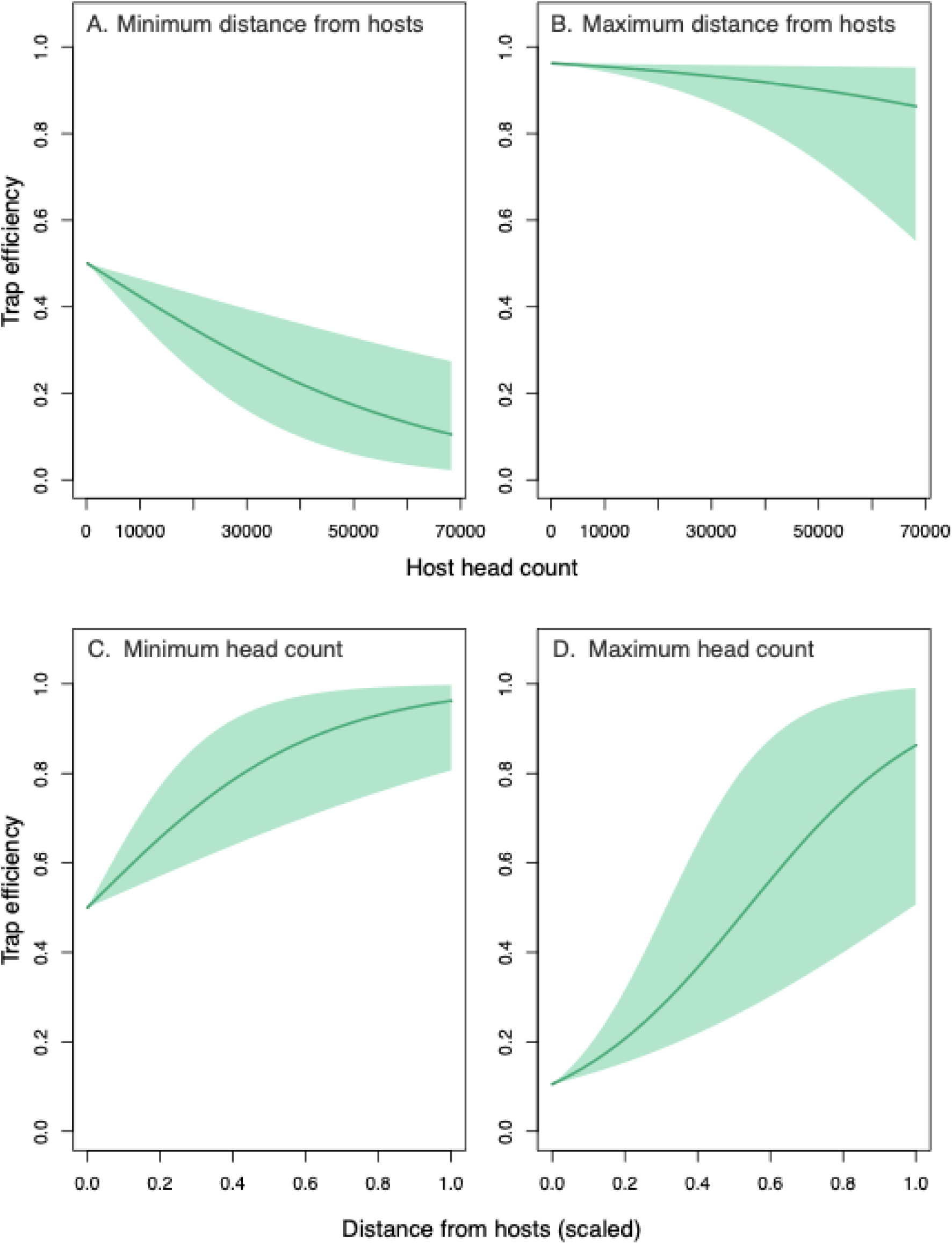
Model predictions of trap efficiency. **A-B:** Estimated effects of *H* on trap efficiency at minimum distance from hosts (panel A) and at maximum distance from hosts (panel B). **C-D:** Estimated effects of *D* on trap efficiency at minimum head count (panel C) and at maximum head count (panel D). Effects of head count *H_s_* and distance *D_s,i_* were calculated using median estimates of all other covariates and their coefficients except for when minimum and maximum were specified. Solid lines are median estimates, and shaded areas represent the 80% credible interval.

#### Midge Abundance λ*_tsi_*

The effect of site type (dairy vs. feedlot) on λ*_t,s,i_* was estimated to be slightly negative (*β_FM_* Median: −0.10; 95% CrI: −1.22, 1.01, compared to 95% prior interval −6.28, 6.20), which translates into a central estimate of 10% more midges at feedlots than at dairies. The credible intervals for this estimate are close to evenly spread around zero, and all models estimate values near zero either positive or negative, meaning that there is strong evidence for an effect of site type on λ*_t,s,i_.* Coefficients for microhabitat variables indicated that, compared to the non-host microhabitat types, there was a strong negative effect of host microhabitat type on λ*_t,s,i_* (Model 5 *β_M1M_*median: −2.27; 95% CrI: −3.72, −0.84).

Distance from host aggregation had a negative effect on per-host midge abundance, with median values of *β_DM_* estimated at 0.84 (95% CrI: −0.75 2.17), indicating a decrease in midge abundance for traps located farther from hosts (Figure 5A). This translates to a median of 14.3% fewer midges with each additional 100 m from the host aggregation (95% CrI −27.9, 21.5).

**Figure 5.**
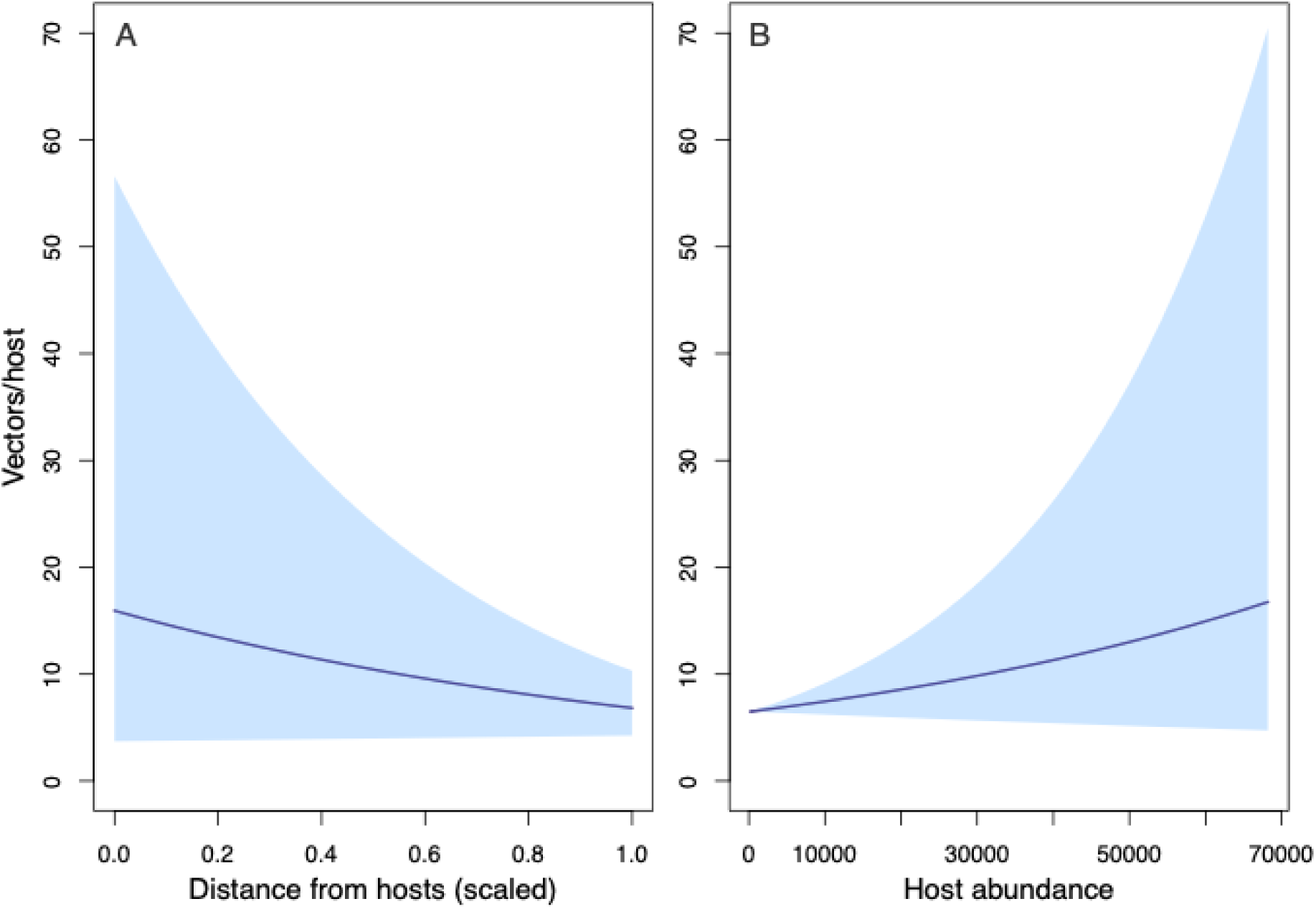
Estimated effects of distance D (panel A) and host abundance H (panel B) on the vector-to-host ratio. Effects of head count D and H were calculated using median estimates of all other covariates and their coefficients. Solid lines are median estimates, and shaded areas represent the 80% credible interval.

Host abundance was estimated to have a positive effect on per-host midge abundance, with a median value of *β_HM_* estimated at 0.95 (95% CrI: −0.96, 3.26) (Figure 5B). Though some estimates of *β_HM_* were below zero, 83% of posterior estimates from this model were greater than zero.

Models assuming a threshold for the effect of host abundance on the vector-to-host ratio (model 5.2 and 5.3 assuming type-II and type-III functional responses, respectively) did not fit the data better than Model 5, which assumed a linear effect of host abundance on vector/host ratio. Though the models with type-II and type-III functional responses had narrower credible ranges for posterior estimates of the effect of host abundance on midge abundance (Figure 6), this was not due to the priors for this relationship being more restrictive than that of the linear effect (Figure S2). LOOIC values for Model 5.2 and Model 5.3 were 0.36 and 0.43 greater than for Model 5, though ΔSE for the three models was greater than ΔLOOIC, so we again cannot definitively choose one model. Perhaps more importantly, the posterior estimates for the parameters describing the shapes of the curves for Models 5.2 and 5.3 are such that the effect of host abundance does not appear to reach a saturation point anyway (Figure 6). Thus, we do not find support for frequency-dependence at high host abundance in this system.

**Figure 6.**
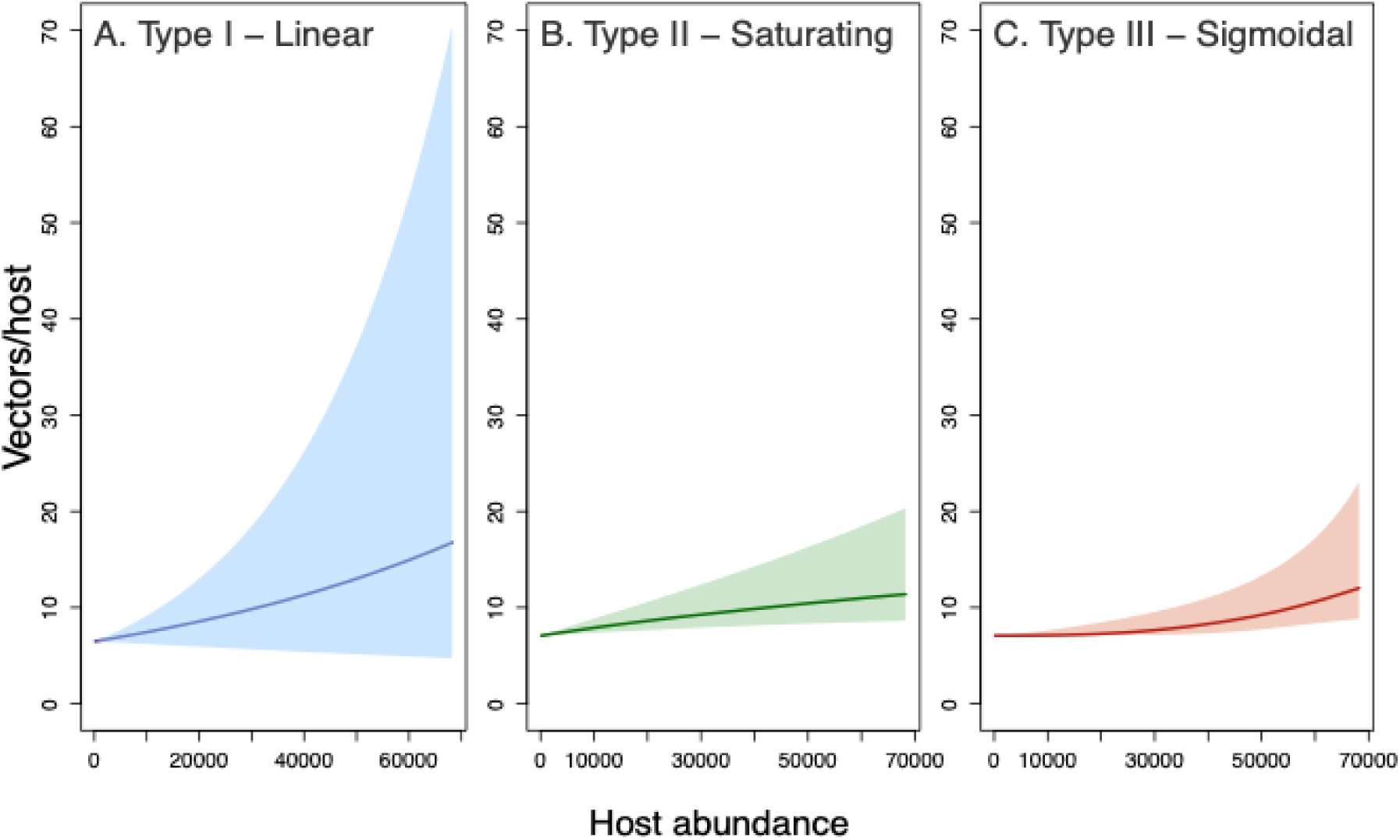
Predicted effects of host abundance *H_s_*on vector/host ratio for models assuming different functional forms. Effect of host abundance *H_s_* was calculated using median estimates of all other covariates and their coefficients. Solid lines are median estimates, and shaded areas represent the areas 80% credible interval. A) Model 5, which assumes a type-I relationship, or linear effect. B) Model 5.2, which assumes a type-II, or saturating relationship. C) Model 5.3, which assumes a type-III or sigmoidal relationship.

### Effect of Host Abundance on Transmission Potential

Given that *R*_0_ is proportional to the vector-to-host ratio, *R*_0_ is expected to vary with host abundance in the same way that *β_HM_* indicates that midge abundance per host does. Our median estimate of *β_HM_* under Model 5 translates into a 2.6-fold increase in *R*_0_ between the site with the lowest host abundance and the site with the highest host abundance (95% CrI: 0.38-25.97). In other words, this is equivalent to a 2.3% increase in the ratio of vectors to hosts for every 1,000 additional hosts. The 95% credibility interval for *β_HM_*suggests, however, that this could be as low as a 0.8% decrease or as high as a 23.2% increase per 1,000 hosts. In addition, we estimated that *R*_0_ should increase as host abundance increases with a probability of 0.83 (Figure 7).

**Figure 7.**
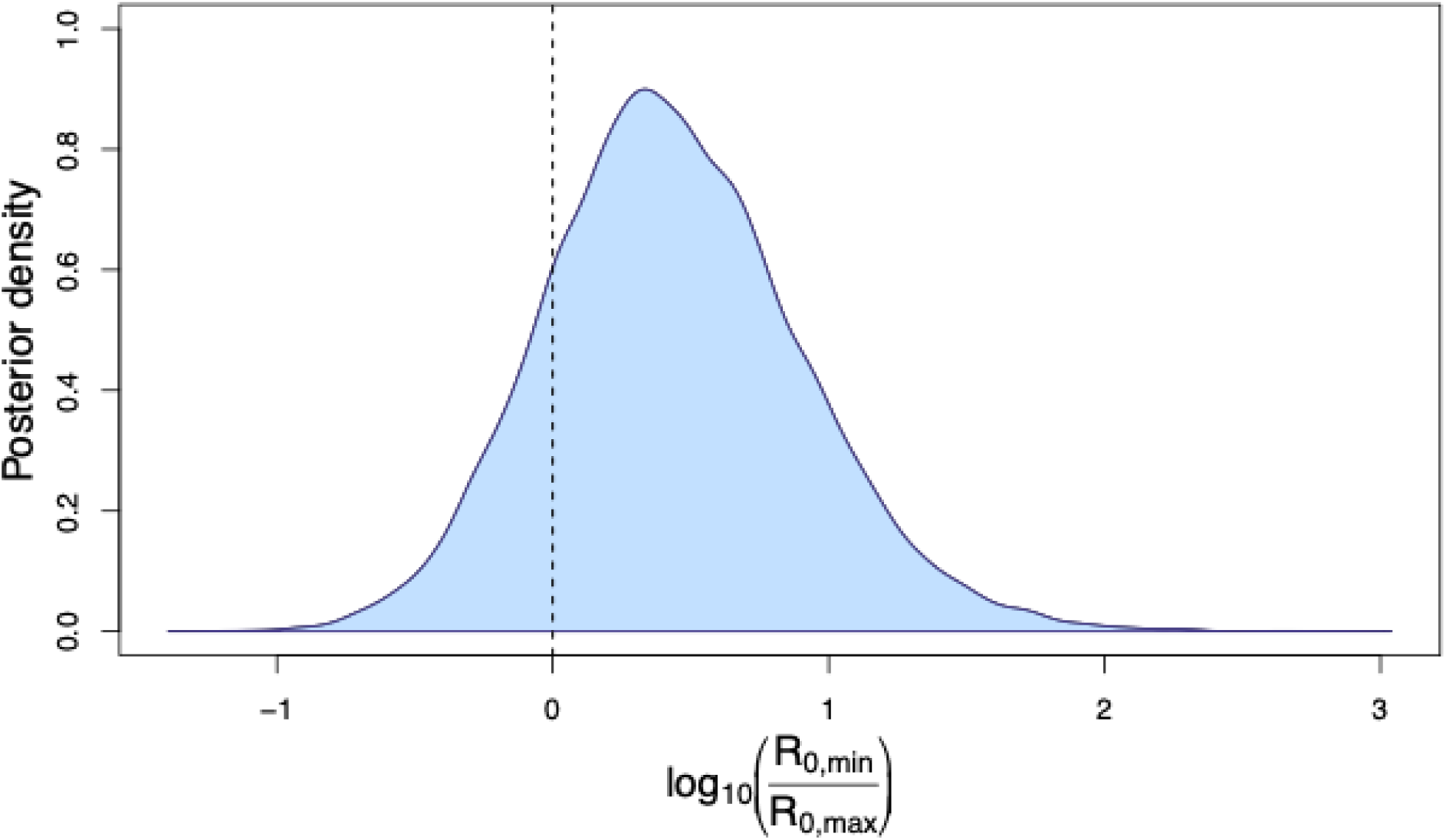
Model 5 posterior density of the predicted ratio of *R*_0_ on the site with highest host abundance to that on the site with lowest host abundance. The dashed line represents *R_0,max_*/*R_0,min_* = 1, because log_10_(1) = 0. Estimates to the right of this line indicate that *R*_0_ increases as host abundance increases.

## Discussion

Overall, our results support an assumption of density-dependent transmission in this system, albeit with non-negligible statistical uncertainty. The parameter *β_HM_*, describing the effect of host abundance on midge abundance per host, was estimated to be positive with probability 0.83. This implies an increase in vectors per host as the number of hosts increases, which is the expected mechanism for density-dependent transmission in a vector-borne pathogen transmission system. Most importantly, this means that the basic reproduction number *R*_0_ - which is directly proportional to the vector-host ratio - should increase as host abundance increases. Therefore, we would expect that BTVs transmission should increase with increased numbers of hosts across farms like the ones that we studied. As a result, farms with more animals should be subject to a higher probability of an outbreak upon pathogen introduction, larger outbreak size, and greater difficulty achieving disease control, consistent with general predictions of pathogen transmission dynamics theory (Keeling et al. 2008). Under an alternative scenario of frequency-dependent transmission, these quantities would be the same on farms with different numbers of animals. Mechanistically, a density-dependent scenario may occur because increased host abundance makes for more blood meals for adult female midges, which would lead to more egg-laying and thus higher midge population. Similarly, more hosts might lead to an increase in breeding habitat, as *Culicoides sonorensis,* in particular, are known for breeding in agricultural wastewater ponds (Purse et al. 2015, Barbera et al. 2025).

Though there is some uncertainty in our results as to whether *β_HM_* is positive, the majority of posterior samples of *β_HM_* were greater than 0 (83% for Model 5, 70 – 85% for all other models). Additionally, median estimates of *β_HM_* remained positive across all models, indicating consistency of our results as other model features changed (Table S2). Interestingly, posterior samples of *β_HM_* below zero correlate with posterior samples of *β_HE_* with lower absolute values. The latter parameter describes the effect of host abundance on trap efficiency, resulting in lower estimates of trap efficiency as a whole. In other words, estimates of the effect of host abundance on midge abundance tended to be negative only when traps were estimated to be capturing midges most efficiently. Given that we believe that, in reality, the traps were not very efficient when placed in close proximity to hosts, we find a positive effect of host abundance on midge abundance per host to be very likely based on the results of our study.

While there was no clear choice of best model based on WAIC and R^2^, the similarity of results among all 12 models indicates that our conclusions are robust to changes in model structure. We chose Model 5 as our primary model to present because we believe that this model is the most biologically justified model. Including an interaction between *D_s,i_* and *H_s_* in the trap efficiency term makes intuitive sense based on our assumption of the trapping mechanism. If a greater number of available hosts detracts from the number of midges attracted by trap CO_2_, one can expect this effect to diminish farther away from the hosts as their CO_2_ dissipates. In contrast, without a clear sense of the spatial scale at which host abundance might affect midge abundance, combined with there not being much change in results when including an interaction in the per-capita midge abundance term, we do not have a strong justification for including that interaction in the final model. With regard to the inclusion of microhabitat variables *M*, models excluding microhabitat predicted a positive effect of distance from hosts on midge abundance (Table S2, panel B), which seems biologically implausible. Models including either binary microhabitat or all microhabitat did not have meaningful differences in parameter estimates (Table S2, panel B) and were not different based on model performance metrics, so we chose the model with binary microhabitat in the interest of reducing unnecessary covariates.

Higher potential for virus transmission by *Culicoides* in areas with high host densities has implications for spread in systems in which transmission is maintained in high-density agricultural clusters. In particular, in the United States, intensively-managed, very high-density operations make up a large portion of the livestock industry, with the number of cows per dairy farm increasing steadily over time as the number of farms has decreased (Barkema et al. 2015, Chase et al. 2006). Commercial feedlot operations have also become larger over time, with 1,000 head or higher feedlots making up 80-85% of the fed cattle market (USDA ERS report). If high-density operations create hotspots for vector abundance, then these environments could function to create hubs of transmission, connected by livestock movement, vector dispersal, or overlap with wildlife populations. In wildlife populations, where host populations may be low, density dependence implies the existence of a threshold host population, below which the pathogen cannot persist (Lloyd-Smith et al. 2005). This allows for the use of culling for disease management, to keep host population size, and thus contact rate, below the threshold (Wobeser 1989).

Although we were able to address the impact of host abundance on vector populations in one setting, there are limitations to our approach. The ratio of vectors to hosts is a theoretically justified proxy for transmission via its relationship with *R*_0_, given that we have no reason to assume that other parameters that affect *R*_0_ should necessarily change with host abundance. However, a measure of infection in the host or the vector would be a more direct and complete way of addressing how much transmission is occurring on different farms. Even so, for pathogens with transmission dynamics that are influenced by acquired immunity among hosts, documenting empirical patterns of infection prevalence that are consistent with density- or frequency-dependent transmission is not straightforward due to nonlinearities between *R*_0_ and infection. Additionally, our study revealed limitations of the exclusive use of CO_2_-baited traps in areas with high host abundance. Use of CO_2_ as an attractant was a justified choice given prior work demonstrating that BTV-infected *Culicoides* are averse to light (McDermott et al. 2015, Mayo et al. 2012, McDermott et al. 2018) and that light traps are less effective in areas with more ambient light. This means that only using light traps could reduce overall trap catch rate, and specifically catch rate of infected midges, resulting in underestimation of vector abundance and pathogen prevalence. However, given our findings that high host abundance interferes with the effectiveness of CO_2_ as an attractant, particularly near the host aggregation, we recommend the use of light traps and CO_2_ traps in combination to obtain more accurate results. Moreover, use of a statistical analysis such as ours - which is capable of estimating true midge abundance as a latent variable based on multiple, imperfect and potentially contradictory data types - would also be advisable.

In conclusion, this study provides an important step in establishing how the transmission of viruses by *Culicoides* scales with cattle density, suggesting that transmission may be magnified in highly concentrated host environments. We also contribute to knowledge of field practice in vector systems that rely on collection of host-seeking vectors, demonstrating the factors that need to be considered when using traps that rely on host-mimicking attractant, while also providing an analytic framework to account for observation processes while elucidating ecological processes. Our findings help to shed light on the role of agricultural practices and human-shaped landscapes in disease maintenance, and can be applied to future work on arboviruses.

## Acknowledgments

This work was funded by USDA-NIFA AFRI grant #2019-67015-28982 as part of the joint USDA-NSF-NIH-BBSRC-BSF Ecology and Evolution of Infectious Diseases program, and USDA-NIFA grant #2021-38420-34065, NSF grant #DEB-2109293, ITE-2333795, BCS-2307944, the Frontier Research Foundation, and the Alliance Bioversity-CIAT.

We gratefully acknowledge producers, staff, and herd managers of the farms from which the samples were obtained.

