## Supplemental info for "Assessing the impact of host density on vector abundance and transmission scaling of *Culicoides*-transmitted pathogens"

**Appendix S1**

*Description of variables and parameters used in the models*

**Table S1.** Descriptions of the variables used in the models, and their prior distributions.

| Variable or Parameter | Description | Prior Distribution |
|---|---|---|
| $\lambda_{tsi}$ | 'True' midge abundance at time *t*, site *s*, and trap location *i*. | - |
| $E_{si}$ | Trap efficiency at site *s* and trap location *i*. | - |
| $H_s$ | Head count at site *s*. | - |
| $V_{tsi}$ | Predicted trapped midge abundance at time t, site s, and trap location i. | - |
| $D_{si}$ | Distance from the host aggregation of trap *i*at site *s*. | - |
| $F_s$ | Site type (dairy or feedlot) of site *s*. | - |
| $RE_{si}$ | Random Effect of trapping location *i* at site *s*. | - |
| $t_t$ | Day in season at trap day *t*. | - |
| $\beta_{HM}$ | Effect of Head Count $H_s$ on true midge abundance $\lambda_{si}$. | Normal(0,2) |
| $\beta_{DM}$ | Effect of distance $D_{si}$ on $\lambda_{si}$. | Normal(0,2) |
| $\beta_{XM}$ | Interacting effect of Head Count $H_s$ and distance $D_{si}$ on $\lambda_{si}$. | Normal(0,2) |
| $\beta_{MIM}$ | Effect of 'Host' habitat type on $\lambda_{si}$. Not included in all models. | Normal(0,2) |

| | | |
|---|---|---|
| $\beta_{M2M}$ | Effect of 'Crop' habitat type on $\lambda_{si}$. Not included in all models. | Normal(0,2) |
| $\beta_{M3M}$ | Effect of 'Lagoon' habitat type on $\lambda_{si}$. Not included in all models. | Normal(0,2) |
| $m_M$ | Percent of distance effect on $\lambda_{si}$ that occurs at the farthest distance. | Uniform(0,1) |
| $\beta_{FM}$ | Effect of Site Type $F_s$ on true midge abundance $\lambda_{si}$. | Normal(0,2) |
| $\beta_{TM}, \tau_1, \tau_2$ | Parameters describing the seasonal curve of midge abundance $\lambda_{si}$. | Normal(0,2), Gamma(3,1), Uniform(0,1) |
| $\sigma_{re}$ | Standard deviation of random effect. | Exponential(1) |
| $\beta_{HE}$ | Effect of Head Count $H_s$ on trap efficiency $E_{si}$. | Gamma(3,1) |
| $\beta_{DE}$ | Effect of distance $D_{si}$ on trap efficiency $E_{si}$. | Gamma(3,1) |
| $\beta_{XE}$ | Interacting effect of Head Count $H_s$ and distance $D_{si}$ on $E_{si}$. | Normal(0,2) |
| $\Phi$ | Variance of $M_{tsi}$. | Gamma(3,1) |
| $a$ | Parameter for Type II and Type III functional forms. | Gamma(2,2) |
| $b$ | Parameter for Type II and Type III functional forms. | Gamma(1,1) |
| $k$ | Parameter for Type III functional form. | Gamma(3,1) |

*Comparison of parameter estimates for all models*

Trap Efficiency $E_{s,i}$

Estimates of the effect of distance on trap efficiency were highest for models that did not include microhabitat variables, with models 1,4,7, and 10 averaging a median estimate of 4.16 for $\beta_{DE}$, compared to 3.28 for the binary-microhabitat models and 3.21 for the models with four microhabitat types (Table S2 N). This difference in model estimates likely reflects collinearity between microhabitat and distance variables, given that the host microhabitat type applies only to traps located at the host aggregation and all non-host microhabitat types apply only to traps located some distance away from it. When no interaction in the efficiency term was included, excluding microhabitat also slightly increased the magnitude of the negative effect of host abundance on trap efficiency (Models 1,7: average median $\beta_{HE}$ = 2.36; model 2,8: average median $\beta_{HE}$ = 2.13; model 3,9: average median $\beta_{HE}$ = 2.12). For models that included that interaction, median estimates of its strength were highest in the models without microhabitat (Model 4,10: average median $\beta_{XE}$ = 0.89) compared to others (model 5, 11: average median $\beta_{XE}$ = 0.71; model 6,12: average median $\beta_{XE}$ = 0.67). In other words, models that did not include microhabitat variables estimated that the negative effect of host abundance on trap function lessened more over greater distances than models including microhabitat, though this did not greatly change estimates of efficiency overall.

Midge Abundance $\lambda_{t,s,i}$

Estimates of the effect of site type, $\beta_{FM,}$ on midge abundance were highest in the models without microhabitat (Models 1, 4, 7, 10 average median $\beta_{FM}$: -0.04) (Table S2E), compared to models with microhabitat included (Models 2, 5, 8, 11 average median $\beta_{FM:}$ -0.10; models 3, 6, 9, 12 average median $\beta_{FM:}$ -0.12).

The strong negative effect of host microhabitat on midge abundance was consistent across models that included microhabitat type (Table S2F). Given that models excluding microhabitat are the only ones that estimate a positive effect of distance on midge abundance under certain conditions, this suggests that the inclusion of microhabitat allows the negative effect seen at host habitat types to be attributed to microhabitat rather than to distance.

For all models that included microhabitat variables as predictors, distance from host aggregation had a negative effect on per-host midge abundance (Models 2, 5, 8, 11 average median $\beta_{DM}$: 0.85; Models 3, 6, 9, 12 average median $\beta_{DM}$: 0.81), indicating a decrease in midge abundance for traps located farther from hosts. Notably, this effect reversed signs in models that excluded microhabitat type (Models 1, 4, 7, 10 average median $\beta_{DM}$: -0.06; This means that models that excluded microhabitat always predict an increase in midges as distance from hosts increases. When models excluding habitat included an interaction between $D$ and $H$ in the $\lambda_{t,s,i,}$ the interaction was estimated to be close to 0 (Model 7,10: median $\beta_{XM}$: -0.06), and did not meaningfully change the effect of distance on vector abundance at different head counts. Other models that include an interaction between $D$ and $H$ in the $\lambda_{t,s,i}$ term estimated a positive interaction (Models 8,11 average median $\beta_{XM}$: 0.19; Models 9,12 average median $\beta_{XM}$: 0.23). Consistent with the data, these models still predict an overall negative effect of distance on observed midge abundance, particularly at high head counts.

Across models, estimates of $\beta_{HM,}$ the parameter describing the effect of host abundance Hs on midge abundance $\lambda_{t,s,i}$, were marginally higher in models that excluded microhabitat (Models 1,4,7,10 $\beta_{HM}$ average median: 1.05; Models 2,5,8,11 $\beta_{HM}$ average median: 0.94; Models 3,6,9,12 $\beta_{HM}$ average median: 0.89).

**Table S2.** Median estimates and 95% credible intervals for all parameters for all models. Columns indicate different parameters, and rows indicate the model number.

| | **A)** $\beta_{HM}$ | | **B)** $\beta_{DM}$ | | **C)** $M_M$ | | **D)** $\beta_{XM}$ | |
|---|---|---|---|---|---|---|---|---|
| Model | Median | 95% CrI | Median | 95% CrI | Median | 95% CrI | Median | 95% CrI |
| 1 | 1.02 | (-1.06, 3.47) | -0.07 | (-1.85, 1.29) | 0.44 | (0.02, 0.97) | NA | NA |
| 2 | 1.01 | (-0.88, 3.29) | 0.85 | (-0.69, 2.17) | 0.49 | (0.03, 0.97) | NA | NA |
| 3 | 0.98 | (-1.03, 3.3) | 0.81 | (-0.86, 2.28) | 0.46 | (0.02, 0.97) | NA | NA |
| 4 | 0.96 | (-1.14, 3.41) | -0.06 | (-1.8, 1.26) | 0.45 | (0.02, 0.97) | NA | NA |
| 5 | 0.95 | (-0.96, 3.26) | 0.85 | (-0.75, 2.17) | 0.48 | (0.02, 0.97) | NA | NA |
| 6 | 0.90 | (-1.11, 3.31) | 0.82 | (-0.77, 2.23) | 0.47 | (0.02, 0.97) | NA | NA |
| 7 | 1.13 | (-1.88, 4.13) | -0.06 | (-1.83, 1.29) | 0.40 | (0.01, 0.97) | -0.07 | (-2.96, 2.74) |
| 8 | 0.91 | (-2.04, 3.87) | 0.86 | (-0.71, 2.22) | 0.45 | (0.02, 0.97) | 0.21 | (-2.59, 3.01) |
| 9 | 0.87 | (-2.1, 3.88) | 0.82 | (-0.79, 2.26) | 0.42 | (0.02, 0.97) | 0.24 | (-2.61, 2.98) |
| 10 | 1.10 | (-1.94, 4.08) | -0.05 | (-1.86, 1.3) | 0.40 | (0.02, 0.96) | -0.06 | (-3.04, 2.86) |
| 11 | 0.90 | (-2.14, 3.85) | 0.85 | (-0.69, 2.21) | 0.45 | (0.02, 0.97) | 0.18 | (-2.62, 2.96) |
| 12 | 0.82 | (-2.25, 3.91) | 0.80 | (-0.83, 2.24) | 0.42 | (0.02, 0.96) | 0.22 | (-2.62, 3.05) |
| 5.2 | NA | NA | 0.87 | (-0.7, 2.16) | 0.5 | (0.03, 0.97) | NA | NA |
| 5.3 | NA | NA | 0.87 | (-0.69, 2.17) | 0.49 | (0.03, 0.97) | NA | NA |

| | E) $\beta_{FM}$ | | F) $\beta_{M1M}$ | | G) $\beta_{M2M}$ | | H) $\beta_{M3M}$ | |
|---|---|---|---|---|---|---|---|---|
| Model | Median | 95% CrI | Median | 95% CrI | Median | 95% CrI | Median | 95% CrI |
| 1 | -0.07 | (-1.32, 1.22) | NA | NA | NA | NA | NA | NA |
| 2 | -0.10 | (-1.23, 1.02) | -2.30 | (-3.7, -0.88) | NA | NA | NA | NA |
| 3 | -0.13 | (-1.3, 1.04) | -2.12 | (-3.75, -0.45) | 0.19 | (-1.26, 1.65) | 0.46 | (-0.93, 1.82) |
| 4 | -0.04 | (-1.27, 1.24) | NA | NA | NA | NA | NA | NA |
| 5 | -0.11 | (-1.22, 1.01) | -2.27 | (-3.72, -0.84) | NA | NA | NA | NA |
| 6 | -0.13 | (-1.29, 1.01) | -2.10 | (-3.74, -0.48) | 0.17 | (-1.27, 1.66) | 0.45 | (-0.92, 1.81) |
| 7 | -0.05 | (-1.32, 1.28) | NA | NA | NA | NA | NA | NA |
| 8 | -0.10 | (-1.23, 1.02) | -2.30 | (-3.72, -0.87) | NA | NA | NA | NA |
| 9 | -0.13 | (-1.28, 1.04) | -2.13 | (-3.76, -0.48) | 0.19 | (-1.24, 1.65) | 0.48 | (-0.92, 1.86) |
| 10 | -0.03 | (-1.33, 1.22) | NA | NA | NA | NA | NA | NA |
| 11 | -0.09 | (-1.21, 1) | -2.29 | (-3.73, -0.86) | NA | NA | NA | NA |
| 12 | -0.12 | (-1.3, 1.06) | -2.12 | (-3.76, -0.47) | 0.19 | (-1.28, 1.65) | 0.44 | (-0.94, 1.85) |
| 5.2 | -0.14 | (-1.23, 0.94) | -2.28 | (-3.67, -0.85) | NA | NA | NA | NA |
| 5.3 | -0.12 | (-1.18, 0.98) | -2.29 | (-3.71, -0.89) | NA | NA | NA | NA |

| | I) $\tau_1$ | | J) $\tau_2$ | | K) $\beta_{tM}$ | | L) $\sigma$ | |
|---|---|---|---|---|---|---|---|---|
| Model | Median | 95% CrI | Median | 95% CrI | Median | 95% CrI | Median | 95% CrI |
| 1 | 1.78 | (0.76, 4.78) | 0.66 | (0.58, 0.84) | 2.22 | (0.73, 4.32) | 1.72 | (1.29, 2.36) |
| 2 | 2.20 | (0.93, 5.52) | 0.66 | (0.58, 0.85) | 1.74 | (0.59, 3.6) | 1.48 | (1.1, 2.06) |
| 3 | 2.34 | (0.94, 5.91) | 0.67 | (0.58, 0.86) | 1.61 | (0.54, 3.53) | 1.53 | (1.14, 2.14) |
| 4 | 1.80 | (0.79, 4.67) | 0.66 | (0.58, 0.83) | 2.20 | (0.74, 4.26) | 1.71 | (1.28, 2.37) |
| 5 | 2.17 | (0.9, 5.44) | 0.66 | (0.58, 0.85) | 1.76 | (0.6, 3.62) | 1.48 | (1.1, 2.05) |
| 6 | 2.34 | (0.94, 5.82) | 0.67 | (0.58, 0.87) | 1.59 | (0.54, 3.53) | 1.53 | (1.13, 2.14) |
| 7 | 1.80 | (0.78, 4.71) | 0.66 | (0.58, 0.84) | 2.20 | (0.75, 4.26) | 1.73 | (1.29, 2.38) |
| 8 | 2.16 | (0.91, 5.43) | 0.66 | (0.58, 0.87) | 1.76 | (0.6, 3.56) | 1.49 | (1.11, 2.05) |
| 9 | 2.36 | (0.96, 5.9) | 0.67 | (0.58, 0.87) | 1.59 | (0.54, 3.48) | 1.54 | (1.14, 2.14) |
| 10 | 1.80 | (0.78, 4.72) | 0.66 | (0.58, 0.84) | 2.19 | (0.74, 4.26) | 1.73 | (1.28, 2.4) |
| 11 | 2.18 | (0.89, 5.46) | 0.66 | (0.58, 0.86) | 1.75 | (0.6, 3.6) | 1.48 | (1.1, 2.04) |
| 12 | 2.30 | (0.95, 5.83) | 0.67 | (0.58, 0.87) | 1.63 | (0.55, 3.51) | 1.54 | (1.14, 2.14) |
| 5.2 | 2.16 | (0.92, 5.37) | 0.66 | (0.58, 0.86) | 1.76 | (0.61, 3.62) | 1.47 | (1.1, 2.03) |
| 5.3 | 2.14 | (0.9, 5.41) | 0.66 | (0.58, 0.87) | 1.77 | (0.59, 3.63) | 1.46 | (1.08, 2.03) |

| | M) $\beta_{HE}$ | | N) $\beta_{DE}$ | | O) $\beta_{XE}$ | | P) $\varphi$ | |
|---|---|---|---|---|---|---|---|---|
| Model | Median | 95% CrI | Median | 95% CrI | Median | 95% CrI | Median | 95% CrI |
| 1 | 2.36 | (0.6, 5.07) | 4.23 | (1.2, 9.4) | NA | NA | 1.66 | (1.15, 2.35) |
| 2 | 2.09 | (0.57, 4.6) | 3.31 | (0.86, 8.13) | NA | NA | 1.65 | (1.14, 2.36) |
| 3 | 2.08 | (0.54, 4.7) | 3.24 | (0.81, 7.92) | NA | NA | 1.65 | (1.15, 2.34) |
| 4 | 2.44 | (0.65, 5.25) | 4.11 | (1.11, 9.48) | 0.90 | (-2.76, 4.51) | 1.65 | (1.15, 2.35) |
| 5 | 2.14 | (0.56, 4.78) | 3.24 | (0.83, 8.26) | 0.74 | (-2.87, 4.32) | 1.65 | (1.14, 2.35) |
| 6 | 2.15 | (0.57, 4.81) | 3.18 | (0.77, 8.05) | 0.68 | (-2.92, 4.34) | 1.65 | (1.15, 2.35) |
| 7 | 2.37 | (0.6, 5.32) | 4.24 | (1.21, 9.4) | NA | NA | 1.65 | (1.15, 2.35) |
| 8 | 2.16 | (0.57, 5.03) | 3.34 | (0.86, 8.01) | NA | NA | 1.65 | (1.15, 2.36) |
| 9 | 2.16 | (0.51, 5.01) | 3.25 | (0.79, 8.07) | NA | NA | 1.65 | (1.14, 2.35) |
| 10 | 2.43 | (0.6, 5.54) | 4.07 | (1.1, 9.36) | 0.89 | (-2.69, 4.57) | 1.65 | (1.14, 2.34) |
| 11 | 2.20 | (0.55, 5.1) | 3.24 | (0.8, 8.17) | 0.69 | (-2.88, 4.31) | 1.65 | (1.15, 2.37) |
| 12 | 2.20 | (0.57, 5.2) | 3.16 | (0.77, 8.04) | 0.67 | (-2.86, 4.29) | 1.64 | (1.14, 2.35) |
| 5.2 | 1.90 | (0.51, 3.95) | 3.35 | (0.85, 8.32) | 0.84 | (-2.73, 4.42) | 1.65 | (1.14, 2.37) |
| 5.3 | 1.92 | (0.54, 4.09) | 3.37 | (0.87, 8.4) | 0.84 | (-2.78, 4.49) | 1.65 | (1.15, 2.37) |

**Table S3**. Median estimates and 95% CrI intervals for the shape parameters for the Hollings type-II and type-III curves, referred to as Model 5.2 and 5.3 because they versions of Model 5, with respective non-linear functions applied to the effect of $H_s$ on $\lambda_{t,s,i}$.

| | **Q)** *a* | | **R)** *b* | | **S)** *k* | |
|---|---|---|---|---|---|---|
| Model | Median | 95% CrI | Median | 95% CrI | Median | 95% CrI |
| 5.2 | 0.84 | (0.13, 2.64) | 0.61 | (0.02, 3.52) | NA | NA |
| 5.3 | 0.95 | (0.15, 2.83) | 0.58 | (0.02, 3.32) | 2,83 | (0.76, 7.42) |

*Supplementary figures*

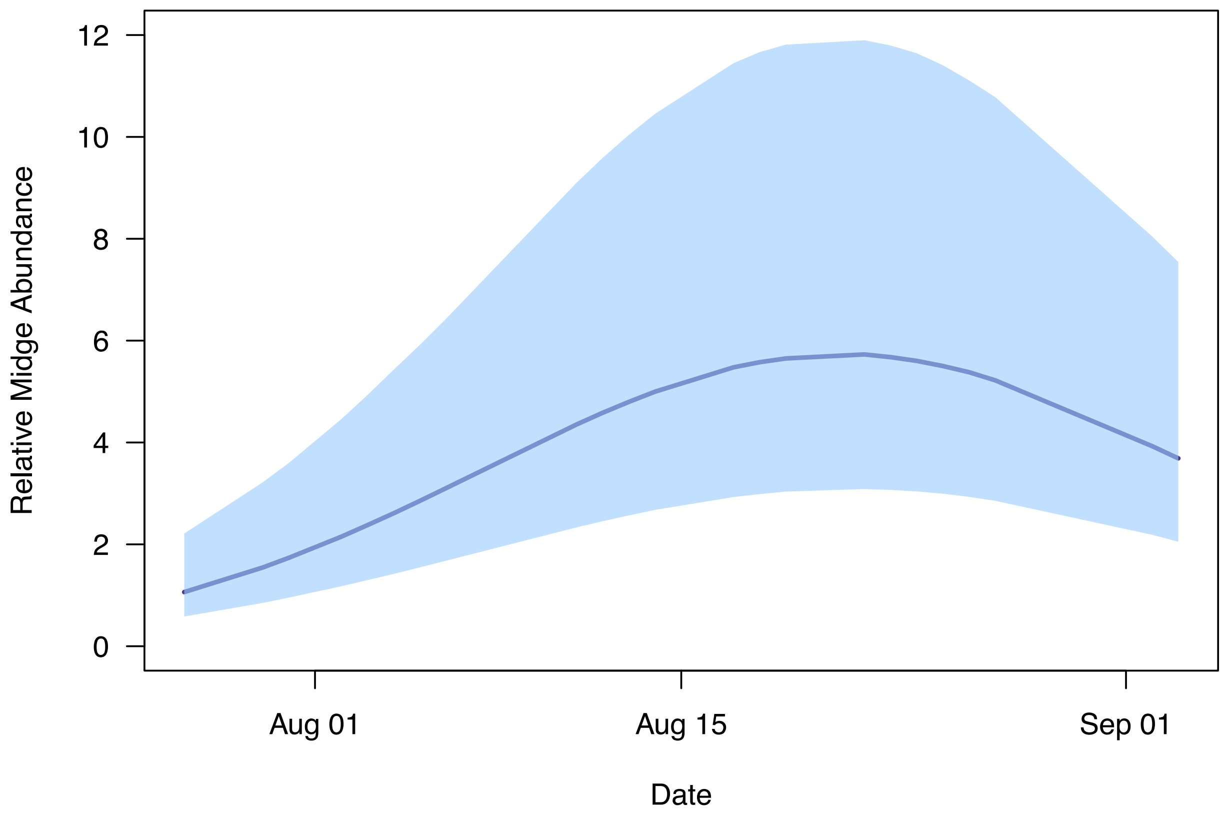


**Figure S1.** Estimated seasonal curve of midge abundance over the course of the collection season. The curve was generated using all posterior samples of $\beta_{tM}$, $\tau_1$, and $\tau_1$ for Model 5. The solid line shows the median curve, and the shaded region the interquartile range.

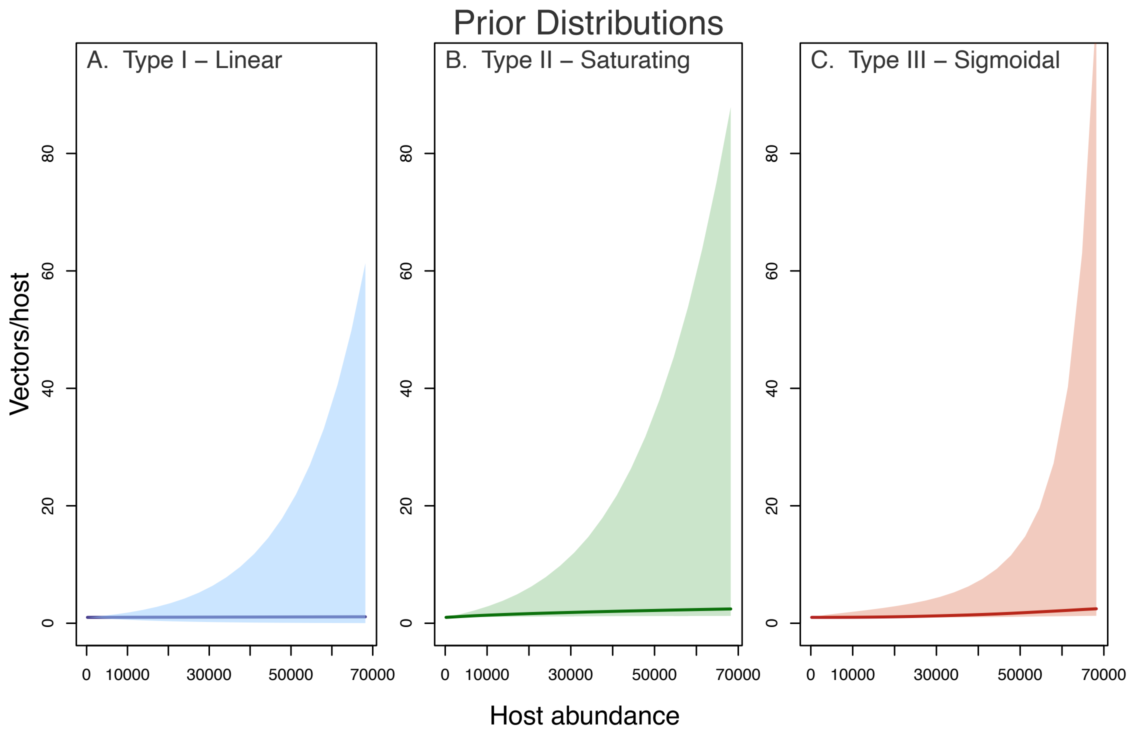


**Figure S2.** Prior distributions for the curves describing the effect of host abundance H on vector/host ratio for models assuming different functional forms. A) Model 5, which assumes a type-I relationship, or linear effect. B) Model 5.2, which assumes a type-II, or saturating relationship. C) Model 5.3, which assumes a type-III or sigmoidal relationship. All models estimate an approximately linear effect of host abundance.
